# Physcraper: a python package for continual update of evolutionary estimates using the Open Tree of Life

**DOI:** 10.1101/2020.09.15.299156

**Authors:** Luna L. Sanchez Reyes, Martha Kandziora, Emily Jane McTavish

## Abstract

1. Phylogenies are a key part of research in many areas of biology. Tools that automate some parts of the process of phylogenetic reconstruction, mainly molecular character matrix assembly, have been developed for the advantage of both specialists in the field of phylogenetics and nonspecialists. However, interpretation of results, comparison with previously available phylogenetic hypotheses, and selection of one phylogeny for downstream analyses and discussion still impose difficulties to one that is not a specialist either on phylogenetic methods or on a particular group of study.
2. Physcraper is a command-line Python program that automates the update of published phylogenies by adding public DNA sequences to underlying alignments of previously published phylogenies. It also provides a framework for straightforward comparison of published phylogenies with their updated versions, by leveraging upon tools from the Open Tree of Life project to link taxonomic information across databases.
3. Physcraper can be used by the nonspecialist, as a tool to generate phylogenetic hypotheses based on publicly available expert phylogenetic knowledge. Phylogeneticists and taxonomic group specialists will find it useful as a tool to facilitate molecular dataset gathering and comparison of alternative phylogenetic hypotheses (topologies).
4. The Physcraper workflow demonstrates the benefits of doing open science for phylogenetics, encouraging researchers to strive for better sharing practices. Physcraper can be used with any OS and is released under an open-source license. Detailed instructions for installation and use are available at https://physcraper.readthedocs.

## 1 Introduction

Phylogenies capture the shared history of organisms and provide key evolutionary context for our biological observations. Public biological databases constitute an amazing resource for evolutionary studies. Updating existing phylogenies with molecular data that has never been incorporated into any phylogenetic estimate, geographical location, fossils, and other data in a reproducible and continuous manner is possible by establishing a data interoperability framework for biological databases. Here, we introduce Physcraper, a tool that automates database connections to build upon homology hypotheses that taxon specialists have assessed and deemed appropriate for a specific phylogenetic scope to update a starting tree and single locus alignments with public DNA data.

Taxonomic idiosyncrasies across databases represent a huge challenge for automatic integration of data into phylogenies, which can be addressed with a unified taxonomy for name standardization. The Open Tree of Life project (OpenTree) constructs a comprehensive tree of life by synthesizing published phylogenies and taxonomy. OpenTree’s “synthetic” tree comprises 2.3 million tips, of which around 90,000 are supported by phylogenies - the remaining 1.4 million taxa are placed in the tree based on taxonomy. To achieve this, OpenTree unifies taxonomic data from various databases (Rees & Cranston 2017), including the USA National Center for Biodiversity Information (NCBI) molecular database GenBank (Benson *et al*. 2000), among others. The OpenTree taxonomy represents a key resource for connecting data from any biological database that has been integrated to it.

Another challenge for incorporating public molecular data into existing phylogenies is assembling high-quality homology hypotheses. While genomics has, and will continue to, revolutionize phylogenetic inference, the variety of alternative genomic sequencing approaches it uses produce largely non-overlapping genomic datasets across taxa, creating challenges in wide scale phylogenetic reconstruction. Phylogenomics ameliorate this problem by focusing on targeted capture of informative loci (Andermann *et al*. 2020). Yet, decades of single locus sequencing have generated massive amounts of homologous DNA datasets that can be used for phylogenetic reconstruction at many scales.

More than a decade ago, GenBank release 159 (April 15, 2007) already hosted 72 million DNA sequences that were gauged to have the potential to resolve phylogenetic relationships of 98.05% of the almost 241,000 distinct taxa in the NCBI taxonomy at the time (Sanderson *et al*. 2008). Assembling a DNA alignment from such a massive database can be done “by hand”, but it is a largely time consuming and mostly non-reproducible approach. Computational pipelines that mine DNA databases fast, efficiently, and reproducibly, have been applied to infer phylogenetic relationships in a variety of organisms (e.g., Smith *et al*. 2009; Izquierdo-Carrasco *et al*. 2014; Antonelli *et al*. 2017). However, fine-grained curated markers and alignments can improve phylogenetic reconstructions, even in phylogenomic analyses (Fragoso-Martínez *et al*. 2017).

There are almost 8,200 publicly available, peer-reviewed alignments, covering around 100,000 distinct taxa in the TreeBASE database (Piel *et al*. 2009), which can be used as seeds to mine molecular databases, and as “jump-start” alignments for phylogenetic reconstructions (Morrison 2006) to continually enrich, update and compare existing phylogenetic knowledge.

Physcraper is a Python pipeline using OpenTree’s taxonomy and programmatic access protocols (API’s) to implement a database interoperability framework that automatically links phylogenies that have been standardized to OpenTree taxonomy, to alignments from TreeBASE, data from GenBank, and phylogenies from OpenTree’s Phylesystem. Physcraper aims to demonstrate the benefits of reproducible workflows and open science in phylogenetics, and encourage better data sharing practices in the community.

## 2 The Physcraper framework

The general Physcraper framework consists of 4 steps (Fig. 1): 1) identifying and processing a tree and its underlying alignment; 2) performing a BLAST search of DNA sequences from original alignment on GenBank, and filtering of new sequences; 3) profile-aligning new sequences to original alignment; 4) performing a phylogenetic analysis and comparing the updated tree to existing phylogenies.

**Figure 1:**
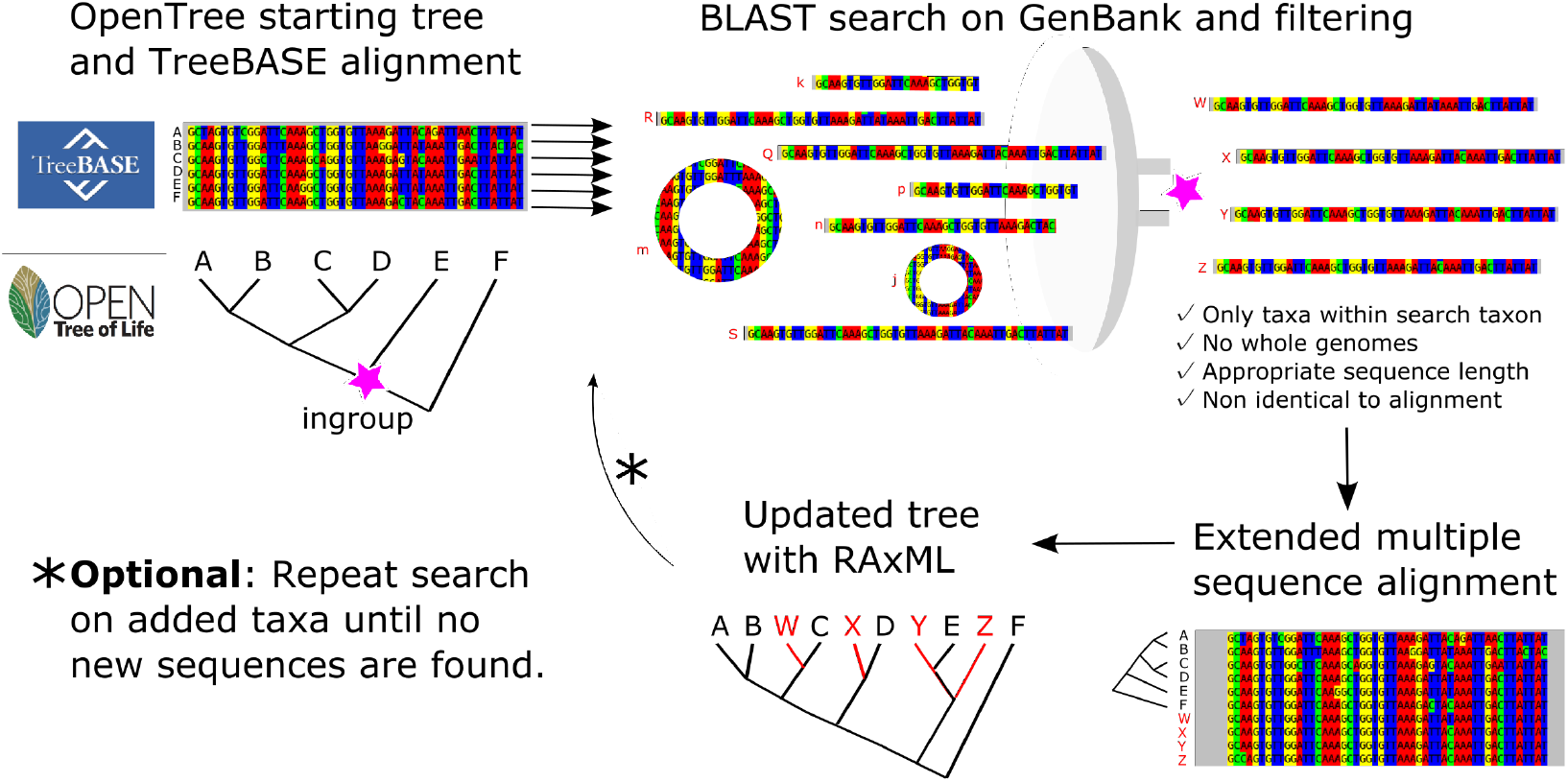
The Physcraper framework consists of 4 steps (see text). The software is fully described on its documentation website at physcraper.readthedocs.io, along with installation instructions, function usage descriptions, examples and tutorials.

### 2.1 The inputs: a tree and an alignment

Taxon names in the input tree must be standardized to OpenTree taxonomy (Rees & Cranston 2017) using OpenTree’s bulk Taxonomic Name Resolution Service TNRS tool. Users can upload their own tree, or choose from among the 2, 950 standardized trees stored in OpenTree’s Phylesystem that also have alignments available on TreeBASE (Piel *et al*. 2009).

The input alignment is a single locus DNA dataset that was used in part or in whole to generate the input tree. Physcraper retrieves TreeBASE alignments automatically. Alternatively, users must provide the path to a local copy of the alignment. Only taxa that are both in the sequence alignment and in the tree are considered further for analysis; at least one taxon and its corresponding sequence are required.

### 2.2 DNA sequence search and filtering

The Basic Local Alignment Search Tool, BLAST (Altschul *et al*. 1990) is used for DNA sequence search on a remote or local GenBank database. It is constrained to a “search taxon”, a taxonomic group in the NCBI taxonomy that is automatically identified using the OpenTree API (Rees & Cranston 2017), as the most recent common ancestor of ingroup taxa that is also a named clade in the NCBI taxonomy (Fig. 1). Alternatively, users can arbitrarily define a search taxon that is either a more or less inclusive clade relative to the ingroup taxa.

BLAST is implemented with the blastn function (Camacho *et al*. 2009) and the BioPython (Cock *et al*. 2009) BLAST function from NCBIWWW module modified to accept an alternative BLAST address. Each sequence in the alignment is BLASTed once against all DNA sequences in GenBank. New sequences are excluded for analysis if they 1) are not in the search taxon; 2) have an e-value above the cutoff (default to 0.00001); 3) fall outside a min and max length threshold, defined as the proportion of the average length without gaps of all sequences in input alignment (default values of 80% and 120%, respectively); 4) or if they are either identical to or shorter than an existing sequence in the input alignment and they represent the same taxon in OpenTree or NCBI taxonomy. An arbitrary maximum number of randomly chosen sequences per taxon are allowed (default to 5).

Reverse, complement, and reverse-complement sequences are identified and translated using BioPython internal functions (Cock *et al*. 2009). Iterative cycles of BLAST searches can be performed, by blasting all new sequences until no new ones are found. By default only one BLAST cycle is performed.

### 2.3 New DNA sequence alignment

MUSCLE (Edgar 2004) is used to perform a profile alignment in which the original alignment is used as a template of homology criteria to align new sequences. The final alignment is not further automatically checked, and additional inspection and refinement are recommended.

### 2.4 Tree reconstruction and comparison

RAxML (Stamatakis 2014) is implemented to reconstruct a Maximum Likelihood (ML) gene tree for each input alignment with default settings (GTRCAT model and 100 bootstrap replicates with default algorithm), using input tree as starting tree for ML searches. Bootstrap results are summarized using DendroPy’s SumTrees module (Sukumaran & Holder 2010).

Physcraper’s main result is an updated phylogenetic hypothesis for the search taxon. Updated and original tree are compared with Robinson-Foulds weighted and unweighted metrics estimated with Dendropy (Sukumaran & Holder 2010), and with a node by node comparison between the synthetic OpenTree and original and updated tree individually, using OpenTree’s conflict API (Redelings & Holder 2017).

## 3 Case Study: The hollies

A user is interested in phylogenetic relationships within the genus *Ilex*. Commonly known as “hollies”, the genus encompasses between 400-700 living species, and is the only extant clade within the family Aquifoliaceae, order Aquifoliales of flowering plants.

An online literature review in June 2020 (Google scholar search for “ilex phylogeny”) reveals that there are several published phylogenies showing relationships within the hollies (Cuénoud *et al*. 2000; Setoguchi & Watanabe 2000; Selbach-Schnadelbach *et al*. 2009; Manen *et al*. 2010), but only two have data publicly available (Gottlieb *et al*. 2005; Yao *et al*. 2020). Gottlieb *et al*. (2005) made original tree and alignment available in TreeBASE. The “Gottlieb2005” tree sampling 41 species was added to OpenTree Phylesystem and it has been integrated into OpenTree’s synthetic tree.

The most recent *Ilex* tree (Yao *et al*. 2020) is available in OpenTree Phylesystem and in the DRYAD repository. With 175 tips, the “Yao2020” tree is the best sampled phylogeny yet available for the hollies.

We ran Physcraper on a laptop Linux computer to update an internal transcribed spacer DNA region (ITS) alignment from Gottlieb *et al*. (2005), using a local GenBank database. BLAST and RAxML analyses ran for 19hrs 45min, with bootstrap analyses taking an additional 13hrs. The updated Gottlieb2005 tree (Fig. 2) displays all 41 distinct taxa from the original study plus 231 new tips, contributing phylogenetic data to 84 additional *Ilex* taxa. The best RaxML tree is 99% resolved, with 25% of nodes with bootstrap support <0.1 and 48% nodes with bootstrap support > 0.75. A large portion of internal branches are negligibly small, with 30 branches < 0.00001 substitution rate units, from which only 9 have a bootstrap support > 0.75 (Fig. 2). For comparison, Yao2020 also contains all 41 distinct taxa from the original Gottlieb2005 study, and contributes phylogenetic data to 134 additional *Ilex* taxa, from which 67 are also in updated Gottlieb2005.

**Figure 2:**
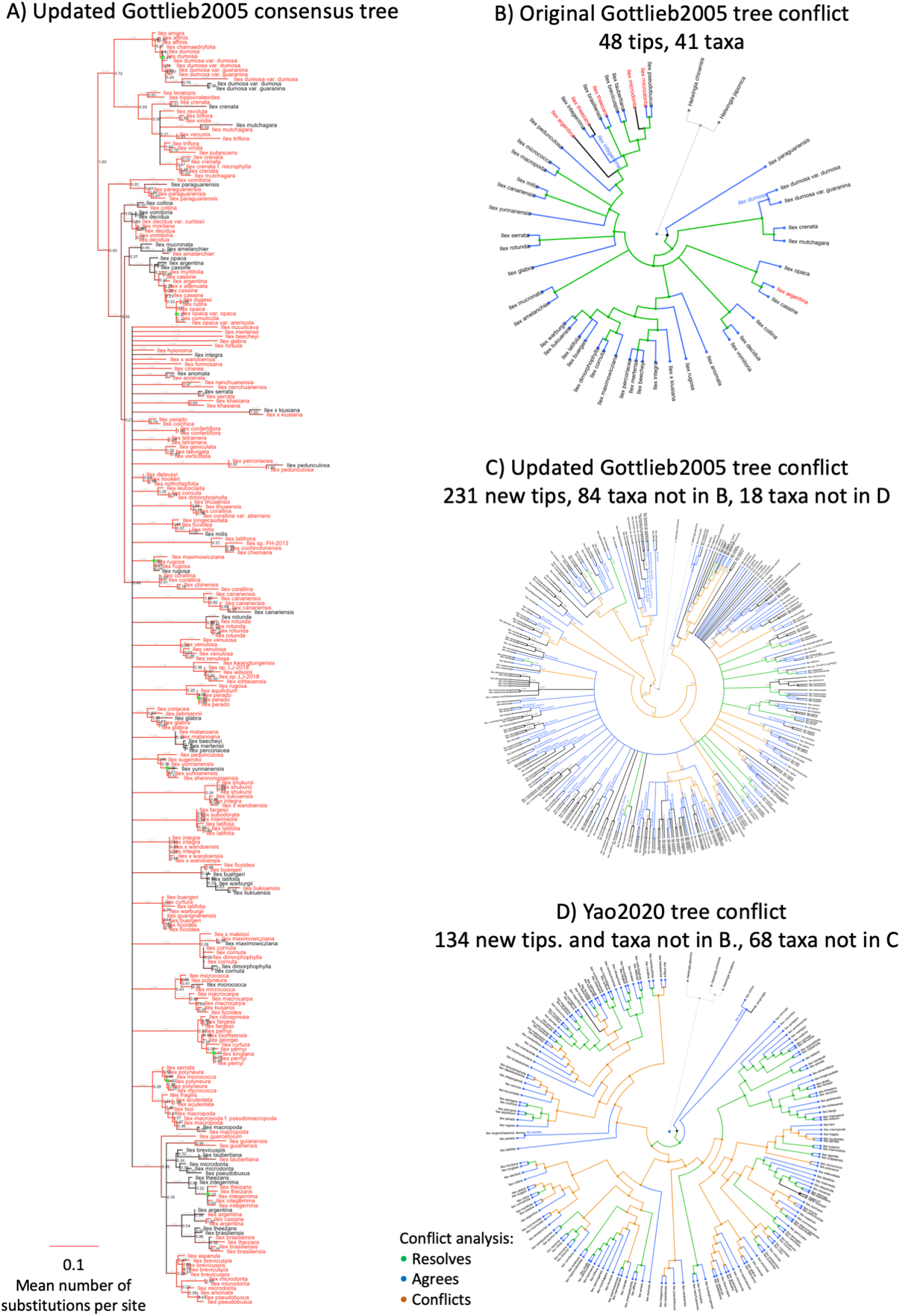
A) Phylogeny updated with Physcraper from Gottlieb et al. 2005 tree in B. Tips in original alignment and new tips added with Physcraper are depicted in black and red, respectively. Physcraper obtained sequences from the GenBank database via local BLAST of all sequences in the original alignment that generated tree in B), filtered them following criteria from section “DNA sequence search and filtering”, aligned them to original alignment using MUSCLE and performed a phylogenetic reconstruction using RAxML with 100 bootstraps. B-D conflict analyses performed with OpenTree tools.

While Yao *et al*. (2020) also used ITS as a marker, their GenBank data is not released yet, so Physcraper was unable to incorporate 68 additional taxa into the analysis. However, Physcraper was able to incorporate 18 taxa that were not in Yao2020.

## 4 Discussion

Databases preserving and democratizing access to biological data have become essential resources for science. New molecular data keep accumulating and tools facilitating its integration into existent evolutionary knowledge are needed.

Phylogenetic pipelines designed to make evolutionary sense of the vast amount of public molecular data (e.g., Phylota (Sanderson *et al*. 2008), PHLAWD (Smith *et al*. 2009), SUPERSMART (Antonelli *et al*. 2017)) focus on generating full phylogenies *de novo*, i.e., inferring phylogenetic relationships from a newly generated homology hypothesis, as opposed to e.g., supertrees, that are generated by summarizing previous phylogenetic estimates. While Physcraper does not generate phylogenies *de novo* in a traditional sense, it successfully generates new phylogenetic knowledge, revealing the importance of open science in facilitating phylogenetic placement of public molecular data and accelerating enrichment and updating of phylogenetic relationships in any region of the tree of life. The PUMPER pipeline (Izquierdo-Carrasco *et al*. 2014) also uses the concept of updating pre-existing alignments to incorporate public molecular data into phylogenies. Unfortunately, installation was unsuccessful following instructions from the author, and a comparison analysis was unfeasible.

Physcraper generates individual gene trees, failing to capture the complexity of species’ evolutionary history (Song *et al*. 2012). Yet, Physcraper facilitates gathering alignments and gene trees for multiple loci from a group of interest, that can be used to reconstruct species trees with ASTRAL (Mirarab *et al*. 2014), BEAST2 (Bouckaert *et al*. 2019), or SVD Quartets (Chifman & Kubatko 2014)).

Physcraper can potentially link phylogenies to data available in any of the taxonomies integrated in the OpenTree taxonomy (Rees & Cranston 2017), such as geographical locations from the Global Biodiversity Information Facility, or fossils from the Paleobiology Database. The Physcraper workflow can be used to rapidly (in a matter of hours) address challenges overarching both fields of ecology and evolution, such as placing newly discovered species phylogenetically (Webb *et al*. 2010), systematizing molecular (and other) databases, i.e., curating taxonomic assignations (San Mauro & Agorreta 2010), and generating custom trees for ecological (Helmus & Ives 2012) and evolutionary downstream analyses (Stoltzfus *et al*. 2013).

Data repositories hold more information than meets the eye. Beyond the main data, they are rich sources of metadata that can be leveraged for the advantage of all areas of biology as well as the advancement of scientific policy and applications. Initial ideas about the data are constantly changed by results from new analyses. Physcraper provides a framework for reproducible phylogenetics that has the potential to consistently provide context for these ideas, highlighting the importance of data sharing and open science in the field, biology and science.

## 5 Acknowledgements

Research was supported by the grant “Sustaining the Open Tree of Life”, NSF ABI No. 1759838, and ABI No. 1759846. Computer time was provided by the Multi-Environment Research Computer for Exploration and Discovery (MERCED) cluster from the University of California, Merced (UCM), supported by the NSF Grant No. ACI-1429783.

We thank the members of the OpenTree development team and the “short bar” Science and Engineering Building 1, UCM, joint lab paper discussion group for valuable comments on this manuscript.

The authors have no conflict of interest to declare.

## 6 Authors’ Contributions

LLSR wrote manuscript, alignment code, documentation, performed analyses and developed examples; MK wrote code for ncbidataparser module, filtering of sequences per OTU and using offline blast searches, wrote documentation and tests; EJM conceived study, wrote most of the code, documentation and tests. All authors contributed to the manuscript and gave final approval for publication.

## 7 Data Archiving

Physcraper source code: https://github.com/McTavishLab/physcraper

Documentation: https://physcraper.readthedocs.io/en/latest/index.html

Examples: https://github.com/McTavishLab/physcraperex

Reproducible manuscript: https://github.com/McTavishLab/physcraper_ms

## References

Altschul, S.F., Gish, W., Miller, W., Myers, E.W. & Lipman, D.J. (1990). Basic local alignment search tool. Journal of Molecular Biology, 215, 403–410.

Andermann, T., Torres Jiménez, M.F., Matos-Maraví, P., Batista, R., Blanco-Pastor, J.L., Gustafsson, A.L.S., Kistler, L., Liberal, I.M., Oxelman, B., Bacon, C.D. & Antonelli, A. (2020). A guide to carrying out a phylogenomic target sequence capture project. Frontiers in Genetics, 10, 1–20.

Antonelli, A., Hettling, H., Condamine, F.L., Vos, K., Nilsson, R.H., Sanderson, M.J., Sauquet, H., Scharn, R., Silvestro, D., Töpel, M. & others. (2017). Toward a self-updating platform for estimating rates of speciation and migration, ages, and relationships of taxa. Systematic Biology, 66, 152–166.

Benson, D.A., Karsch-Mizrachi, I., Lipman, D.J., Ostell, J., Rapp, B.A. & Wheeler, D.L. (2000). GenBank. Nucleic Acids Research, 28, 15–18.

Bouckaert, R., Vaughan, T.G., Barido-Sottani, J., Duchêne, S., Fourment, M., Gavryushkina, A., Heled, J., Jones, G., Kühnert, D., Maio, N.D., Matschiner, M., Mendes, F.K., Müller, N.F., Ogilvie, H.A., Plessis, L. du, Popinga, A., Rambaut, A., Rasmussen, D., Siveroni, I., Suchard, M.A., Wu, C.-H., Xie, D., Zhang, C., Stadler, T. & Drummond, A.J. (2019). BEAST 2.5: An advanced software platform for Bayesian evolutionary analysis. PLOS Computational Biology, 15, e1006650.

Camacho, C., George, C., Vahram, A., Ning, M., Jason, P., Kevin, B. & Thomas, L. (2009). BLAST+: Architecture and applications. BMC Bioinformatics, 10, 421.

Chifman, J. & Kubatko, L. (2014). Quartet inference from SNP data under the coalescent model. Bioinformatics, 30, 3317–3324.

Cock, P.J., Antao, T., Chang, J.T., Chapman, B.A., Cox, C.J., Dalke, A., Friedberg, I., Hamelryck, T., Kauff, F., Wilczynski, B. & others. (2009). Biopython: freely available Python tools for computational molecular biology and bioinformatics. Bioinformatics, 25, 1422–1423.

Cuénoud, P., Martinez, M.A. del P., Loizeay, P.-A., Spichiger, R., Andrews, S. & Manen, J.-F. (2000). Molecular phylogeny and biogeography of the genus *Ilex* L.(Aquifoliaceae). Annals of Botany, 85, 111–122.

Edgar, R.C. (2004). MUSCLE: Multiple sequence alignment with high accuracy and high throughput. Nucleic Acids Research, 32, 1792–1797.

Fragoso-Martínez, I., Salazar, G.A., Martínez-Gordillo, M., Magallón, S., Sánchez-Reyes, L., Lemmon, E.M., Lemmon, A.R., Sazatornil, F. & Mendoza, C.G. (2017). A pilot study applying the plant Anchored Hybrid Enrichment method to New World sages (*Salvia* subgenus Calosphace; Lamiaceae). Molecular Phylogenetics and Evolution, 117, 124–134.

Gottlieb, A.M., Giberti, G.C. & Poggio, L. (2005). Molecular analyses of the genus *Ilex* (Aquifoliaceae) in southern south america, evidence from aflp and its sequence data. American Journal of Botany, 92, 352–369.

Helmus, M.R. & Ives, A.R. (2012). Phylogenetic diversity–area curves. Ecology, 93, S31–S43.

Izquierdo-Carrasco, F., Cazes, J., Smith, S.A. & Stamatakis, A. (2014). PUmPER: Phylogenies updated perpetually. Bioinformatics, 30, 1476–1477.

Manen, J.-F., Barriera, G., Loizeau, P.-A. & Naciri, Y. (2010). The history of extant *Ilex* species (Aquifoliaceae): evidence of hybridization within a Miocene radiation. Molecular Phylogenetics and Evolution, 57, 961–977.

Mirarab, S., Reaz, R., Bayzid, M.S., Zimmermann, T., Swenson, M.S. & Warnow, T. (2014). ASTRAL: Genome-scale coalescent-based species tree estimation. Bioinformatics, 30, i541–i548.

Morrison, D.A. (2006). Multiple sequence alignment for phylogenetic purposes. Australian Systematic Botany, 19, 479–539.

Piel, W., Chan, L., Dominus, M., Ruan, J., Vos, R. & Tannen, V. (2009). Treebase v. 2: A database of phylogenetic knowledge. E-biosphere.

Redelings, B.D. & Holder, M.T. (2017). A supertree pipeline for summarizing phylogenetic and taxonomic information for millions of species. PeerJ, 5, e3058.

Rees, J.A. & Cranston, K. (2017). Automated assembly of a reference taxonomy for phylogenetic data synthesis. Biodiversity Data Journal.

Sanderson, M.J., Boss, D., Chen, D., Cranston, K.A. & Wehe, A. (2008). The PhyLoTA Browser: Processing GenBank for Molecular Phylogenetics Research. Systematic Biology, 57, 335–346.

San Mauro, D. & Agorreta, A. (2010). Molecular systematics: A synthesis of the common methods and the state of knowledge. Cellular & Molecular Biology Letters, 15, 311.

Selbach-Schnadelbach, A., Cavalli, S.S., Manen, J.-F., Coelho, G.C. & De Souza-Chies, T.T. (2009). New information for *Ilex* phylogenetics based on the plastid psbA-trnH intergenic spacer (Aquifoliaceae). Botanical Journal of the Linnean Society, 159, 182–193.

Setoguchi, H. & Watanabe, I. (2000). Intersectional gene flow between insular endemics of *Ilex* (Aquifoliaceae) on the Bonin Islands and the Ryukyu Islands. American Journal of Botany, 87, 793–810.

Smith, S.A., Beaulieu, J.M. & Donoghue, M.J. (2009). Mega-phylogeny approach for comparative biology: An alternative to supertree and supermatrix approaches. BMC Evolutionary Biology, 9, 37.

Song, S., Liu, L., Edwards, S.V. & Wu, S. (2012). Resolving conflict in eutherian mammal phylogeny using phylogenomics and the multispecies coalescent model. Proceedings of the National Academy of Sciences, 109, 14942–14947.

Stamatakis, A. (2014). RAxML version 8: A tool for phylogenetic analysis and post-analysis of large phylogenies. Bioinformatics, 30, 1312–1313.

Stoltzfus, A., Lapp, H., Matasci, N., Deus, H., Sidlauskas, B., Zmasek, C.M., Vaidya, G., Pontelli, E., Cranston, K., Vos, R. & others. (2013). Phylotastic! Making tree-of-life knowledge accessible, reusable and convenient. BMC Bioinformatics, 14, 158.

Sukumaran, J. & Holder, M.T. (2010). DendroPy: a Python library for phylogenetic computing. Bioinformatics, 26, 1569–1571.

Webb, C.O., Slik, J.F. & Triono, T. (2010). Biodiversity inventory and informatics in Southeast Asia. Biodiversity and Conservation, 19, 955–972.

Yao, X., Song, Y., Yang, J.-B., Tan, Y.-H. & Corlett, R.T. (2020). Phylogeny and biogeography of the hollies (*Ilex* L., Aquifoliaceae). Journal of Systematics and Evolution, 58, 1–10.

